# SOD1^A4V^ aggregation alters ubiquitin homeostasis in a cell model of ALS

**DOI:** 10.1101/166165

**Authors:** Natalie E. Farrawell, Isabella Lambert-Smith, Kristen Mitchell, Jessie McKenna, Luke McAlary, Prajwal Ciryam, Kara L. Vine, Darren N. Saunders, Justin J. Yerbury

## Abstract

Amyotrophic lateral sclerosis (ALS) is a fatal neurodegenerative disease involving the selective death of upper and lower motor neurons in the primary motor cortex and spinal cord. A hallmark of ALS pathology is the accumulation of ubiquitinated protein inclusions within motor neurons. Previous studies suggest the sequestration of ubiquitin (Ub) into inclusions reduces the availability of free Ub, which is essential for cellular function and survival. However, the dynamics of the Ub landscape in ALS have not yet been described. Here we show that Ub homeostasis is altered in a SOD1 cell model of ALS. Utilising fluorescently tagged Ub, we followed the distribution of Ub in living cells expressing SOD1 and show that Ub is present at the earliest stages of SOD1 aggregation. We also report that cells containing aggregates of mutant SOD1 have greater ubiquitin-proteasome system (UPS) dysfunction as measured by the accumulation of the fluorescent proteasome reporter tdTomatoCL1. Furthermore, SOD1 aggregation is associated with the redistribution of Ub and depletion of the free Ub pool. Ubiquitomics analysis indicates that mutant SOD1 is associated with a shift of Ub to a pool of supersaturated proteins including those associated with oxidative phosphorylation and metabolism, corresponding with altered mitochondrial morphology and function. Taken together, these results suggest misfolded SOD1 contributes to UPS dysfunction and that Ub homeostasis is an important target for monitoring pathological changes in ALS.

## Background

Amyotrophic lateral sclerosis (ALS, also known as motor neuron disease, MND) is a progressive neurodegenerative disease leading to paralysis of voluntary muscles due to the death of motor neurons in the brain and spinal cord. The prognosis of ALS is poor, usually leading to death within 2 to 5 years of first symptoms. A fraction of patients also develop clinical or subclinical frontotemporal dementia (FTD) [1]. In most cases of ALS, the cause remains unknown (sporadic ALS; sALS). However, approximately 10% of cases are inherited (familial ALS; fALS). A large proportion (20%) of fALS cases can be attributed to mutations in the gene encoding superoxide dismutase 1 (SOD1) [2]. SOD1 was the first gene discovered to cause fALS and is also the most widely studied. There are now over 20 genes known to cause ALS [2] including a growing list of genes associated with dysregulation of ubiquitin (Ub) signalling. Genetic mutations in VCP, SQSTM1, UBQLN2, OPTN have all been associated with ALS, and are all part of the cell’s protein degradation machinery. In addition, recently discovered mutations in TBK1 [3] and CCNF [4] add to this growing list of degradation machinery associated with ALS. The precise role of each of these genes in the homeostasis of Ub is unknown, but Ub sequestration into insoluble inclusions is common to all forms of ALS [5].

Abnormal accumulation of proteins into insoluble aggregates is a hallmark of many neurodegenerative diseases, including Alzheimer’s disease, Parkinson’s disease Huntington’s disease and ALS [6]. In the context of ALS, there is growing evidence that a correlation exists between protein aggregate load and neuronal loss in ALS spinal cord [7–11]. Our previous work showed a correlation between *in vitro* aggregation propensity and rate of disease progression [12], suggesting that protein aggregates are intimately linked with motor neuron cell death. Recent work also indicates that protein misfolding and aggregation may be responsible for disease progression through a prion like propagation throughout the nervous system [11, 13-16]. It is unlikely that misfolding alone is responsible for the disease and that post translational modifications play an important role [17]. One crucial post translational modification is ubiquitination, necessary for protein degradation. Degradation defects that lead to inclusion formation are associated with a tendency for cells to be dysfunctional and undergo apoptosis [18–20].

Inclusions associated with neurodegeneration consist of a variety of proteins including proteins specific to the disease (e.g. Aβ and tau in Alzheimer’s disease [21]), proteins associated with cellular quality control machinery (e.g. molecular chaperones [22, 23] and the proteasome [24]) and other unrelated aggregation prone proteins [25, 26]. Based on analysis of human tissue, it has been shown that a large number of proteins are supersaturated in the cell, with cellular concentrations under wild-type conditions that exceed their predicted solubility [25, 26]. These supersaturated proteins are associated with the biochemical pathways underpinning a variety of neurodegenerative diseases. Most recently, we have shown that proteins co-aggregating with SOD1, TDP-43, and FUS inclusions are supersaturated [5], consistent with a collapse of motor neuron protein homeostasis in ALS. Others have found that the proteins that co-aggregate with c9orf72 dipeptide repeats in cell models are also supersaturated [27]. The composition of inclusions found in ALS varies considerably depending on whether the disease is sporadic or familial, and the genetics of the familial forms.

Ub is a pervasive feature of inclusions in ALS, regardless of underlying genetic aetiology. Ub is a versatile signalling molecule responsible for controlling an array of cellular pathways including transcription, translation, vehicle transport and apoptosis [28]. Ub labels substrate proteins via a highly ordered multi-step enzymatic cascade with specific differences in the length and topology of poly-ubiquitin chains determining a range of signalling outcomes, including proteolytic degradation via the proteasome [29, 30]. Inside cells, Ub exists in a dynamic equilibrium between free Ub and Ub conjugates and is controlled by the opposing actions of Ub ligases and deubiquitinating enzymes (DUBS) [31, 32]. It has been proposed that the sequestration of Ub into insoluble aggregates may deplete the free Ub pool required by many essential cellular processes [32].

Although mounting genetic and functional evidence suggests an important role for the UPS in the development of ALS pathology, the distribution and availability of Ub in ALS models has not yet been described. In the work reported here, we sought to characterise the Ub landscape in a cell-based SOD1 model of ALS **(Figure 1)**. We show that Ub homeostasis is disrupted in cells containing SOD1 aggregates by following the distribution of fluorescently labelled Ub in live cells expressing SOD1. The aggregation of mutant SOD1 leads to an accumulation of the proteasome reporter tdTomato^CL1^, indicative of UPS dysfunction. This dysfunction was further supported by the redistribution of the Ub pool and decrease in free Ub levels observed in cells with SOD1 aggregates. Moreover, ubiquitome analysis confirmed misfolded SOD1 was associated with Ub redistribution and subsequent alterations to mitochondrial morphology. This report highlights that disruption to Ub homeostasis is associated with aggregation of misfolded proteins and may play an important role in the pathogenesis of ALS.

**Figure 1:**
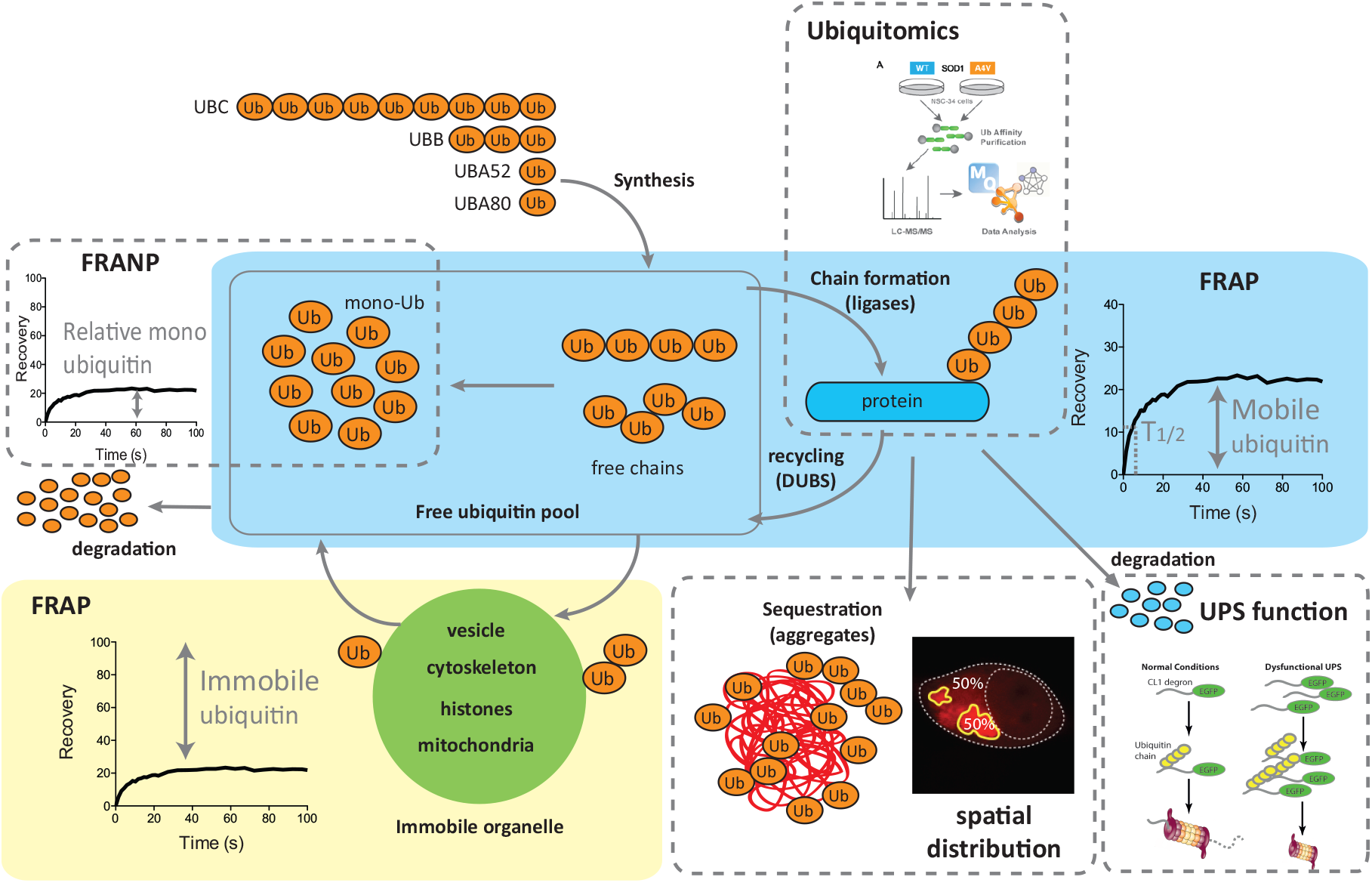
Ubiquitin exists in a dynamic equilibrium in the cell, partitioning into mobile and immobile fractions. Schematic representation of Ub homeostasis, showing the different methods used to interrogate various aspects of this system in this study. Ub is synthesised from several genes, generating a pool of free Ub that consists of monomeric and polymeric chains of Ub. Ub can be conjugated with substrate proteins via the enzymatic activity of a large number of Ub ligases. The cellular fate or outcome of such ubiquitylation is dictated by the chain type formed by Ub. The mobile fraction of Ub measured by fluorescence recovery after photobleaching (FRAP) is the sum of monomeric and polymeric free Ub and diffusible ubiquitylated protein (blue box). The immobile pool of Ub is that conjugated to large slowly diffusing structures such as vesicles, organelles, histones and cytoskeleton (green box). Ubiquitin proteasome system (UPS) degradation function can be compared using fluorescent reporters. Generally accumulation of reporters suggests increased competition, or decreased rates of degradation. The ubiquitin-modified proteome can be quantified using enrichment capture and mass spectrometry (ubiqitomics). A subset of mobile Ub is free monomeric Ub can be measured using fluorescence recovery after nucleus photobleaching (FRANP). Monomeric Ub is the only form of tagged Ub that can diffuse through the nuclear pore, so by bleaching the entire nucleus we can measure monomeric Ub as it diffuses back in. In the context of protein aggregation a proportion of Ub is conjugated to substrates within the inclusions, we can quantify the proportion of Ub incorporated into inclusions using image analysis.

## Methods

### Plasmids

pEGFP-N1 vectors containing human SOD1^WT^ and SOD1^A4V^ were generated as described [33]. GFP-Ub and mRFP-Ub (Addgene plasmids 11928 and 11935, provided by Nico Dantuma [31, 34]) were acquired from Addgene (USA). The mcherry-Ub construct was created by replacing the mRFP sequence in mRFP-Ub with mcherry fluorescent protein. The tdTomato^CL1^ construct was obtained by cloning the CL1 sequence (ACKNWFSSLSHFVIHL) into pcDNA3.1(+)tdTomato.

### Cell Culture and Transfection

Neuroblastoma × Spinal cord hybrid NSC-34 cells [35] were maintained in Dulbecco’s Modified Eagles Medium/Ham’s Nutrient Mixture F12 (DMEM/F12) supplemented with 10% foetal bovine serum (FBS, Bovogen Biologicals, Australia). Cells were maintained at 37 °C in a humidified incubator with 5% atmospheric CO_2_. For confocal microscopy, cells were grown on 13 mm round coverslips in 24-well plates or on 8-well µ-slides (Ibidi, Germany). Cells were grown in 6-well plates for cell lysate and cell sorting experiments. Cells were transfected using Lipofectamine 3000 (Invitrogen, USA) according to manufacturer’s instructions with 0.5 μg DNA per well for a 24-well plate, 0.2 μg DNA per well for 8-well µ-slides and 2.5 μg DNA per well for 6-well plates. For co-transfections the amount of DNA was divided equally between constructs.

### Measurement of UPS function

To quantify UPS activity, NSC-34 cells were co-transfected with the fluorescent proteasome reporter tdTomato^CL1^ and treated overnight (~18 h) with 0-30µM of the proteasome inhibitor MG132. The level of tdTomato^CL1^ fluorescence was determined using flow cytometry on a Becton Dickinson flow cytometer 48 h post transfection as in [36].

### Live cell imaging

Imaging of NSC-34 cells co-transfected with SOD1^A4V^-GFP and mRFP-Ub was performed 24 h post transfection on a Leica TCS SP5 confocal microscope. Cells were imaged every 15 min over 17 h in a CO_2_ chamber maintained at 37°C with 5% CO_2_. For fluorescence recovery after photobleaching (FRAP) and fluorescence recovery after nucleus photobleaching (FRANP) experiments, analysis was performed on transfected NSC-34 cells 48 h post transfection using the LAS AF FRAP Application Wizard on the Leica TCS SP5 confocal Microscope. Images were acquired using the 63× objective with two line averages and a scan speed of 700 Hz. Five pre-bleach images were acquired over 7.5 s with the 561 nm laser set at 20% power. The region of interest (ROI) was then bleached using the ‘zoom in ROI’ method over five frames of 1.5 s at 100% laser power. For FRANP analysis, the entire nucleus was bleached. Fluorescence recovery or loss was monitored for 120 s with the laser power set back at 20%.

### Frequency distribution analysis

GFP fluorescence in NSC-34 cells expressing soluble and insoluble (aggregated) SOD1^A4V^-GFP was quantified from confocal images taken 48 h post transfection using ImageJ 1.48v [37]. Frequency distribution analysis was subsequently performed in GraphPad Prism version 5.00 for Windows (GraphPad software, USA).

### Immunofluorescence

NSC-34 cells grown on coverslips were transfected with GFP-tagged mutant SOD1 and fixed for 20 min at room temperature (RT) with 4% paraformaldehyde (PFA) (Merck Millipore, USA) in phosphate buffered saline (PBS) 48 h post transfection. Cells were permeabilized in 1% tritonX-100 (TX-100) in PBS for 30 min on ice before blocking for 1 h at RT with 5% FBS, 1% Bovine serum albumin (BSA) 0.3% TX-100 in PBS. Cells were incubated with rabbit primary antibodies against K48 or K63 Ub chain linkages (05-1307/05-1308, Merck Millipore) overnight at 4 °C followed by Alexa Fluor 647-conjugated anti-rabbit IgG secondary antibody for 5 h at RT. All antibodies were diluted in 1% BSA, 0.1% TX-100 in PBS and cells were washed with PBS between each incubation step.

Spinal cord sections from the SOD1^G93A^ mouse were also stained for SOD1 and K48 or K63 Ub chain linkages. A pap pen (Daido Sangyo, Tokyo, Japan) was used to separate tissue sections mounted onto the same slide. Sections were then fixed with 4% PFA for 15 min at RT before permeabilization at RT for 10 min with 0.1% TX-100 in PBS containing 2% (v/v) normal horse serum (NHS). Sections were then blocked for 20 min at RT with 20% NHS and 2% BSA in PBS followed by staining overnight at 4 °C in a humidified chamber with sheep anti-superoxide dismustase-1 (ab8866, Abcam, UK), mouse neuron-specific beta III tubulin antibody (ab78078, Abcam) and rabbit anti-ubiquitin K48 or K63 (Merck, Millipore). The following day, sections were incubated with Alexa Fluor-conjugated secondary antibodies reactive to sheep, mouse and rabbit IgG for 1 h at RT. Following staining, coverslips were mounted onto slides using ProLong Diamond Antifade Mountant (Molecular probes, USA). All imaging experiments were analysed on a Leica TCS SP5 confocal microscope.

### Cell lysis

NSC-34 cells grown in 6-well plates and transfected with SOD1-GFP or mRFP-Ub were harvested 48 h post transfection with trypsin/EDTA (Gibco, USA). Cells were washed with PBS before being resuspended in RIPA buffer (50 mM TrisHCl pH 7.4, 1% (w/v) sodium deoxycholate, 150 mM NaCl, 1 mM EDTA, 1% TX-100, 0.1% SDS, 10mM NEM, 1 mM sodium orthovanadate, Halt^TM^ Protease Inhibitor Cocktail (Thermo Scientific, USA)). Protein concentration was determined by BCA assay.

### Western blot

Cell lysates with a total protein concentration of 100 µg were reduced with β-mercaptoethanol and heated for 10 min at 70 °C before being analysed on a Mini-PROTEAN TGX Stain-Free Gel (Bio-Rad, USA) for 2 h at 100 V. Proteins were transferred onto a nitrocellulose membrane (Pall Corporation, USA) using the standard protocol on the Bio-Rad Trans-Blot Turbo Transfer System. The membrane was blocked in 5% skim milk powder in Tris buffered saline, 0.2% (v/v) Tween-20 (TBST) for 1 h at RT before probing for Ub with rabbit anti-ubiquitin antibody (ab137025, Abcam) overnight at 4 °C. The following day the membrane was washed three times with TBST over 30 min before incubating with horseradish peroxidise-conjugated anti-rabbit antibody (1706515, Bio-Rad) for 1 h at RT. The membrane was visualised with chemiluminescent substrate (Thermo Scientific) on the Amersham Imager 6600RGB.

### Cell sorting

NSC-34 cells transfected with SOD1-GFP were harvested 48 h post-transfection with trypsin-EDTA (Gibco) and washed with PBS before being resuspended in resuspension buffer (PBS containing 25 mM HEPES, 1 mM EDTA and 1% FBS) and filtered through a 40 µm nylon mesh strainer (Greiner Bio-One, Austria). Enrichment of GFP positive cells was performed on a Bio-Rad S3e cell sorter. Cells were sorted based on GFP fluorescence (excitation 488 nm, emission 525/30nm) and collected into 5 mL FACS tubes containing DMEM/F12 supplemented with 10% FBS and 2 × penicillin/streptomycin (Gibco). Following enrichment, cells were lysed in RIPA buffer (as above) for ubiquitomics analysis.

### Ubiquitomics

Proteomics to identify ubiquitin-modified proteome (ubiquitome) in NSC-34 cells was performed as described in [38]. Cells were lysed in lysis buffer (50 mM Tris-HCl (pH 7.5), 150 mM NaCl, 0.5% NP-40, 1 mM DTT, 1× EDTA-free protease inhibitor cocktail, 10 mM N-ethylmaleimide, 1 mM sodium orthovanadate). Protein concentration was determined using a BCA assay and 300 µg of total protein was used per sample to immunopurify mono- and poly-ubiquitinated proteins using 30 µL of VIVAbind Ub affinity matrix (Ubiquitin Kit, VIVA Bioscience). Samples were incubated for 2 h at 4 °C with end-to-end mixing then matrix was collected by centrifugation (1 min; 4 °C; 1000 *×* g) and washed six times in detergent free wash buffer (500 µL). After washing, beads were digested for 30 min at 27 °C, then reduced with 1 mM DTT and left to digest overnight at room temperature with sequencing-grade trypsin (5 µg/mL, Promega), as described previously [39]. Samples were alkylated with 5 mg/mL iodoacetamide and protease digestion terminated with trifluoroacetic acid. Trypsinized eluents were collected after brief centrifugation then purified and desalted using self-packed tips with 6 layers of C18 Empore disks (Pacific Laboratory Products), then dried in a SpeedVac. Samples were then resuspended in 12 µL 5% formic acid, 2% acetonitrile and stored at -80 °C.

For nLC-MS/MS analysis, 5 uL of each of the peptide samples were loaded and separated along a C18 column (400 mm, 75 µm ID, 3 µm silica beads) and introduced by nanoelectrospray into an LTQ Orbitrap Velos Pro coupled to an Easy-nLC HPLC (Thermo Fisher). Tandem mass spectrometry data was collected for the top 10 most abundant ions per scan over a 140-minute time gradient. The order of data collection was randomized to interchange between biological conditions with BSA run between each sample to minimize temporal bias. MS/MS raw files were analyzed using the Andromeda search engine integrated into MaxQuant (v1.2.7.4) [40] against the Uniprot Mouse database. A false discovery rate of 1% was tolerated for protein, peptide, and sites, and one missed cleavage was allowed. MaxQuant output data were filtered to remove contaminants, reverse hits, proteins only identified by site and proteins with < 2 unique peptides [41]. An individual protein was defined as present under a particular condition if it was detected in a minimum of two replicates. Protein-protein interactions and KEGG pathway analysis among the resulting protein list was analyzed using STRING (v10) [42] (with a confidence score of 0.700) and Cytoscape (v3.1.1) [43].

### Analysis of Mitochondrial Morphology and Function

NSC-34 cells transiently transfected with SOD1-GFP were incubated with 100nM Mitotracker Deep Red FM (Molecular probes) or Mitotracker CMXRos (Molecular probes) for 30 min at 37 °C prior to analysis by confocal microscopy or flow cytometry 48 h post transfection. Mitochondrial morphology was examined using a mitochondrial morphology macro [44] on ImageJ 1.48v. Mitotracker Red and Mitotracker CMXRos accumulation in cells expressing high levels of SOD1-GFP was also assessed by flow cytometry (Becton Dickinson).

## Results

### SOD1 aggregates are ubiquitylated early

We previously showed that all cellular SOD1 aggregates contained Ub at the time points tested [45]. Here we used real-time imaging using a Ub fusion protein to follow inclusion formation in a single cell from its genesis until it had taken up a large proportion of the cytoplasm. Previous work has demonstrated that the behaviour of GFP-Ub fusions in cells is indistinguishable from endogenous Ub and provide a robust fluorescent indicator of Ub distribution [31]. We created a mCherry-Ub fusion protein that behaved in a similar manner to the GFP fusion protein (supplementary **Figure S1**). Ub was observed in foci at the earliest detectable stages of SOD1 aggregation (**Figure 2A**, see insert). Ub was continuously added to inclusions throughout their formation, as was SOD1 (**Figure 2A**). To ensure that the Ub accumulation in SOD1 inclusions was not an artefact of Ub over expression, we next probed for endogenous Ub using antibodies. SOD1^A4V^ expression causes inclusions in approximately 15% of NSC-34 cells (**Figure S2)** and in those cells that contain inclusions we observed a high degree of overlap between SOD1 aggregates and Ub when using both K48 and K63 chain specific antibodies (**Figure 2B**). We also found that SOD1 aggregates contained K48 chains in SOD1^G93A^ mouse spinal motor neurons (**Figure 2C**).

**Figure 2:**
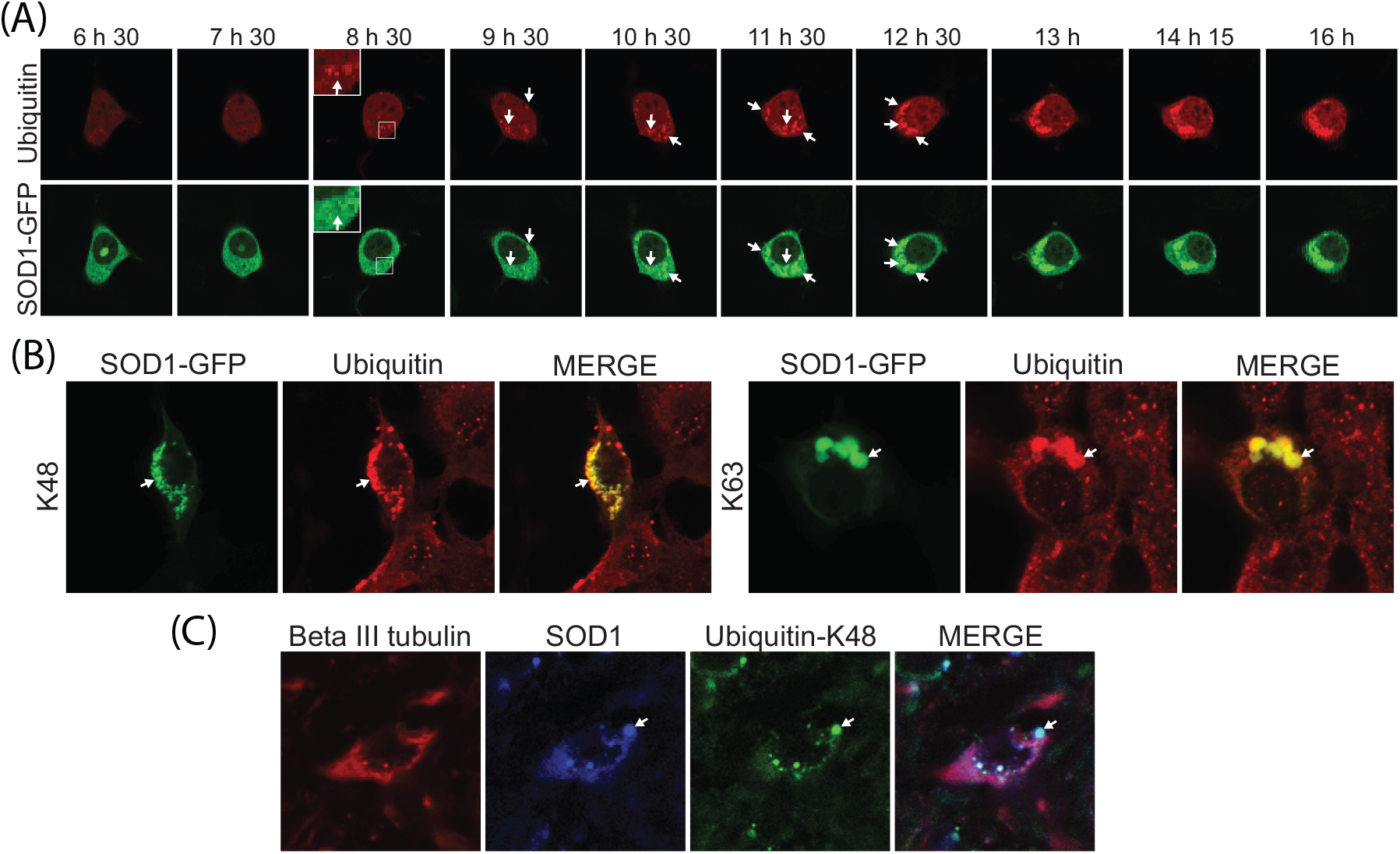
SOD1 co-aggregates with ubiquitin. (A) NSC-34 cells co-transfected with SOD1^A4V^-GFP and mRFP-Ub were imaged every 15 min for 17 h. (B) NSC-34 cells overexpressing SOD1^A4V^-GFP were fixed, permeabilized and stained for Ub_K48_ and Ub_K63_ polyubiquitin chains 48 h post transfection. (C) Spinal cord from the SOD1^G93A^ mouse was fixed, permeabilized and stained for neuron-specific beta III tubulin, SOD1 and Ub_K48_. Co-localization of SOD1 and Ub is indicated with arrows.

### Cells with SOD1 aggregates have increased UPS dysfunction

We previously showed that a UPS reporter containing a CL1 degron peptide for rapid degradation accumulates to significantly higher amounts in ALS patient fibroblasts compared to controls, suggesting an overwhelmed UPS [36]. Work from others suggests that SOD1 aggregates are toxic because they interfere with the quality control function of the juxtanuclear quality control compartment (JUNQ) [20]. To examine the relationship between SOD1 aggregation and UPS dysfunction we used transiently transfected NSC-34 cells that contained SOD1^A4V^-GFP and tdTomato^CL1^. We initially used flow cytometry to examine the effect of proteasome inhibition on reporter accumulation. While there was a significant difference in reporter signal between SOD1 and SOD1^A4V^ expressing cells, both cell lines showed a similar dose-dependent increase in reporter signal with MG132 treatment (**Figure 3A**). This suggests that, while SOD1^A4V^ expression causes significant UPS disruption, there is no specific vulnerability to proteasome inhibition of cells expressing mutant SOD1 compared to wild-type expressing cells.

**Figure 3:**
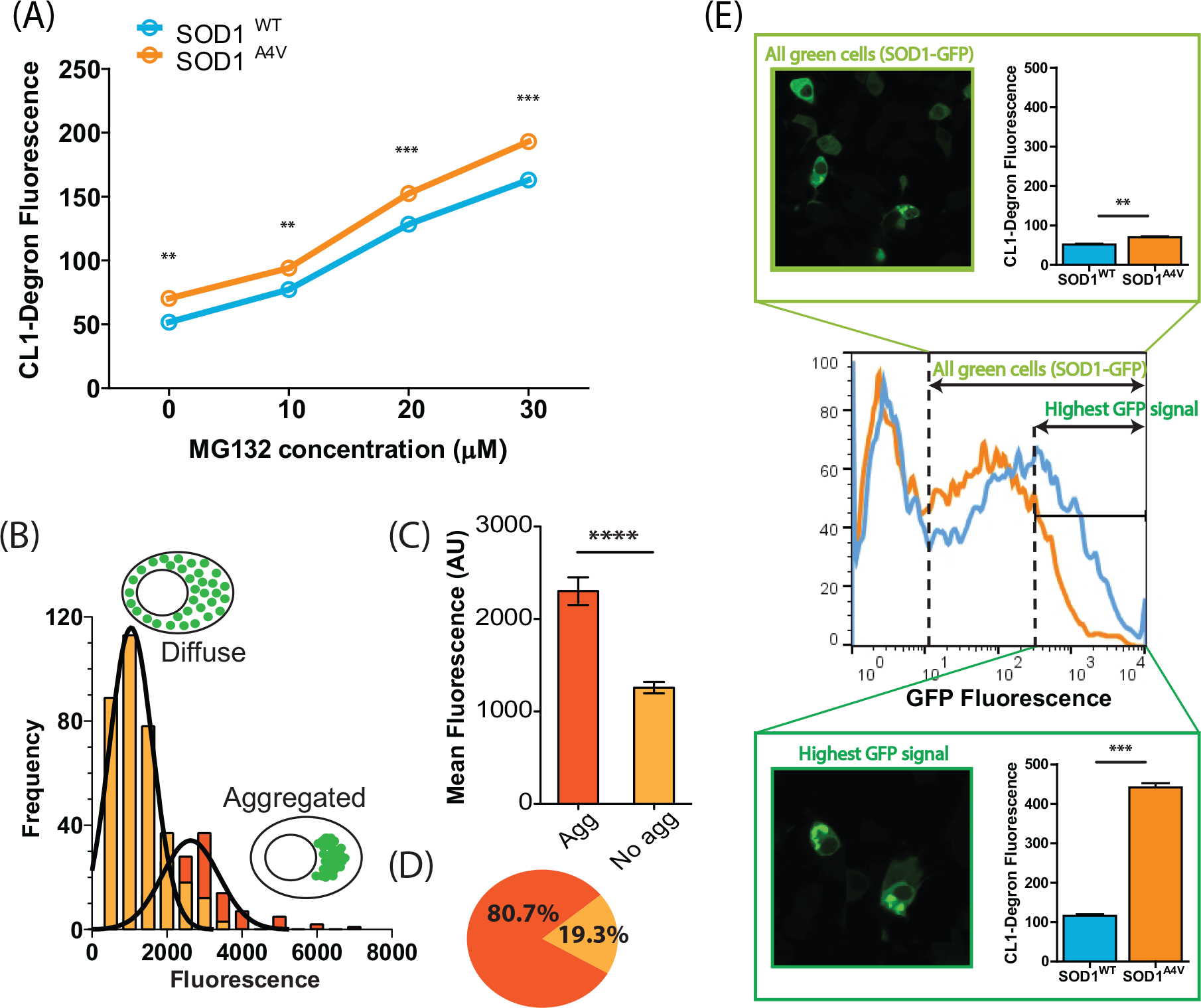
Mutant SOD1^A4V^ alters UPS activity. (A) Dose dependant response of UPS activity (tdTomato^CL1^ fluorescence) in NSC-34 cells co-transfected with SOD1-GFP after overnight treatment with the proteasome inhibitor MG132. Data represent mean CL1-Degron fluorescence ± SEM (n=3), ** P< 0.01; *** P<0.001. (B) Frequency distribution analysis of SOD1-GFP fluorescence was performed on cells expressing soluble (diffuse) SOD1^A4V^ and aggregated SOD1^A4V^. (C) Cells containing SOD1^A4V^ aggregates exhibited significantly higher fluorescence than cells that did not contain aggregates, **** P< 0.0001. (D) 80% of cells expressing the highest SOD1-GFP signal (top 10%) contained aggregates. (E) Cells expressing the highest GFP signal typically contained aggregates and revealed greater significant differences in tdTomato^CL1^ fluorescence between cells expressing SOD1^WT^ and SOD1^A4V^.

However, this analysis examined the entire population of SOD1^WT^ or SOD1^A4V^ expressing cells and previous work using a similar CL1-containing fluorescent reporter suggests that the aggregation of Huntingtin and CFTR cause UPS dysfunction [46]. We hypothesized that inclusions formed by mutant SOD1 might influence UPS activity in a similar fashion, a phenomenon that may not be observed when analysing the entire population of cells, given that only 15% of NSC-34 cells expressing SOD1^A4V^ produce inclusions (**Figure S2**). In order to identify cells with SOD1 inclusions in our flow cytometry data we tested the relationship between aggregation and total cellular fluorescence (**Figure 3B**). Cells were transiently transfected with SOD1^A4V^-GFP and imaged 48 h post transfection using confocal microscopy. The total cellular fluorescence for individual cells was plotted for both cells with and without inclusions. The fluorescence of both populations was normally distributed (**Figure 3B**). The mean fluorescence of cells containing aggregates was significantly greater than that of cells without aggregates (**Figure 3C**), presumably due to continued accumulation of SOD1 into inclusions. However, there was significant overlap between the two populations so we could not distinguish populations entirely. We decided to analyze the most highly fluorescent cells (top 10%) across the entire population. In this subset of cells, approximately 80% contained SOD1^A4V^ inclusions (**Figure 3D**). We reasoned that analyzing the top 10% fluorescent cells would sufficiently enrich for cells containing inclusions. Using flow cytometry, we observed a 4× increase in reporter fluorescence (P< 0.001) - indicating UPS dysfunction - in the top 10% of SOD1^A4V^–GFP expressing cells compared to control cells in the same fluorescence range. That is, UPS dysfunction in SOD1^A4V^ cells is largely restricted to cells containing aggregates. By comparison, only a small increase (1.2×) in reporter fluorescence was observed in the entire population of SOD1^A4V^–GFP expressing cells compared with the entire SOD1^WT^ -GFP expressing population (P< 0.01) (**Figure 3E**). This suggests that SOD1 aggregation is associated with compromised UPS function.

### Ubiquitin pools are disturbed in cells containing SOD1 aggregates

Ub exists in a dynamic equilibrium in the cell, partitioning into four major pools; i) immobile in the nucleus, ii) immobile in the cytoplasm, iii) soluble polyUb chains and iv) a small fraction as free monomeric Ub [31]. Mobile Ub is a combination of free monomeric Ub, free Ub chains and Ub attached to diffusible proteins (**Figure 1**). Immobile Ub is primarily bound to histones in the nucleus and bound to organelles and cytoskeleton in the cytoplasm (**Figure 1**). To determine if the observed UPS dysfunction in cells with mutant SOD1 aggregates was associated with altered Ub homeostasis we examined the relative distribution of Ub into different cellular pools following expression of wild-type or mutant SOD1 genes by FRAP (**Figure 4A-E**). The diffusion rate back into the bleached area provides information on the mobility of the Ub species present in solution and the amount of immobile fluorophore in the compartment of interest (i.e. bound to membrane vesicles or cytoskeleton in the cytosol, and bound to histones in the nucleus). We selected ROIs in both the nucleus and cytoplasm (**Figure 4B**) in cells co-expressing mCherry-Ub and either GFP, SOD1^WT^-GFP or SOD^A4V^-GFP. In the case of SOD^A4V^-GFP expressing cells, we performed the analysis on two subpopulations, with and without aggregates. As previously reported, we find that Ub fluorescence recovery after bleaching (indicating Ub mobility) is higher in the cytoplasm than in the nucleus, likely due to a large proportion of nuclear Ub being attached to histones. Patterns of recovery appeared similar in all treatments, with the exception of a lower cytoplasmic recovery in the cells containing SOD^A4V^-GFP aggregates (**Figure 4A**). After calculating the mean half-life (T_1/2_) of recovery in each case, we find no significant differences between any of the cell populations (**Figure 4D**). This suggests that within the mobile population of Ub, the presence of SOD^A4V^-GFP aggregates does not significantly alter the kinetics of Ub diffusion.

**Figure 4:**
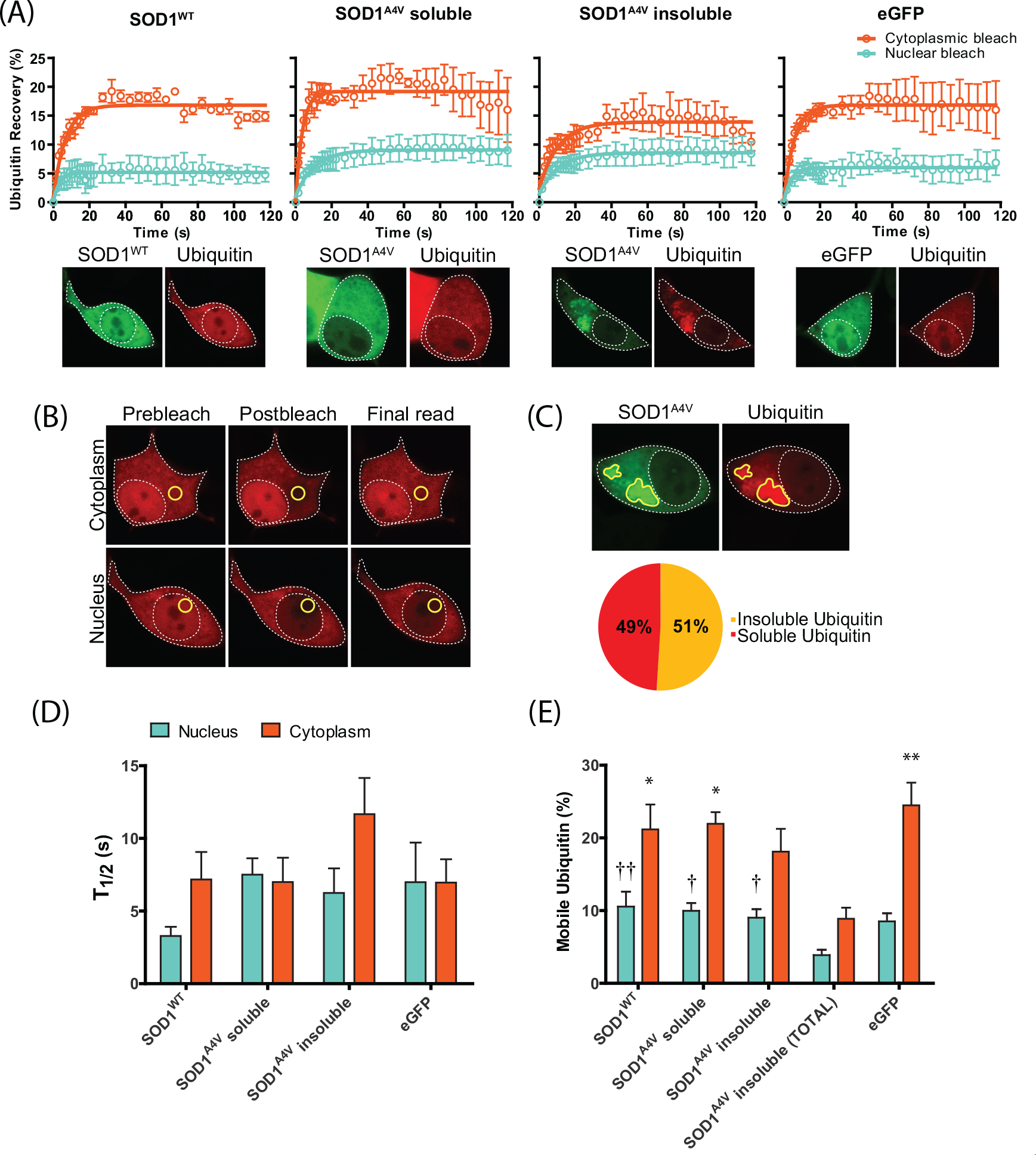
Ubiquitin distribution is altered in cells containing SOD1^A4V^ aggregates. (A) NSC-34 cells co-transfected with SOD1-GFP and mCherry-Ub were photobleached in either the nucleus or cytoplasm and recovery of Ub fluorescence was monitored for 120 s. Data shown are means ± SEM (n=3) and are representative of 3 independent experiments. (B) Representative confocal images of pre-bleach, post-bleach and recovery endpoint are shown with the ROI marked in yellow. (C) The amount of Ub incorporated into SOD1^A4V^ inclusions was quantified using ImageJ. (D) Diffusion rates of mCherry-Ub measured in both the nucleus and cytoplasm of co-transfected NSC-34 cells. Error bars are SEM (n ≥ 6, combined from 3 independent experiments). (E) Quantification of the proportion of mobile Ub in the nucleus and cytoplasm of cells expressing either SOD1^WT^, soluble SOD1^A4V^ or insoluble SOD1^A4V^. SOD1^A4V^ insoluble (TOTAL) represents total Ub levels taking into account the proportion of Ub incorporated into SOD1 inclusions (determined in (C)). Data shown are means ± SEM combined from 3 independent experiments (n ≥ 7). One way Anova was used to compare differences. * and † indicate significant difference to SOD1^A4V^ insoluble (TOTAL) in the cytoplasm and nucleus respectively. * † P < 0.05, ** † † P < 0.01.

This observation prompted us to investigate whether SOD^A4V^-GFP aggregates alter the amount of mobile Ub available in the cell. We find that on average, 51% of the cellular Ub signal is detected in aggregates, indicating that the presence of insoluble aggregates alters the distribution of the Ub pool (**Figure 4C**). The significance of this aggregate-associated fraction of Ub is clear in the context of our analysis of the relative distribution of mobile Ub pools. Based on our FRAP analysis, which does not consider the aggregate-associated Ub, there is no significant difference in the mobility of Ub between SOD^A4V^-GFP cells and controls (**Figure 4E**). However, if we correct for the 51% of Ub we know to be immobile in cells with inclusions, we find that the actual mobile Ub pool in SOD^A4V^-GFP cells is significantly smaller than in controls (**Figure 4E**).

### Free monomeric ubiquitin is lowered in cells with SOD1 aggregates

To test whether the depletion of mobile Ub in cells containing SOD^A4V^-GFP aggregates may diminish free monomeric Ub, we next examined the relative amounts of monomeric Ub in cells expressing GFP, SOD1^WT^-GFP, or SOD^A4V^-GFP for 48 h, by Western blot (**Figure 5A**). We did not observe any significant differences using this method, likely due to the fact that this analysis represents a total measure of all cells in the culture, including non-transfected cells. We therefore turned to fluorescence recovery after nucleus photobleaching (FRANP) analysis to monitor free Ub in single cells. Monomeric mCherry-Ub will diffuse through the nuclear pore (< 60 kDa), while mCherry-Ub incorporated into Ub chains will not [31]. Hence, relative free monomeric Ub can be measured by monitoring diffusion through the nuclear pore following fluorescence bleaching solely in the nucleus or cytosol [31]. We transfected cells with GFP, SOD1^WT^-GFP, or SOD^A4V^-GFP (**Figure 5B**) for 48 h, then bleached the entire nucleus and quantified the amount of relative monomeric Ub diffusing through the nuclear pore back into the nucleus (**Figure 5C**). We observed a significant drop in free monomeric Ub in cells containing SOD^A4V^-GFP aggregates, but no difference in the level of monomeric Ub between cells expressing SOD1^WT^ and SOD^A4V^ in the absence of inclusions (**Figure 5D**).

**Figure 5:**
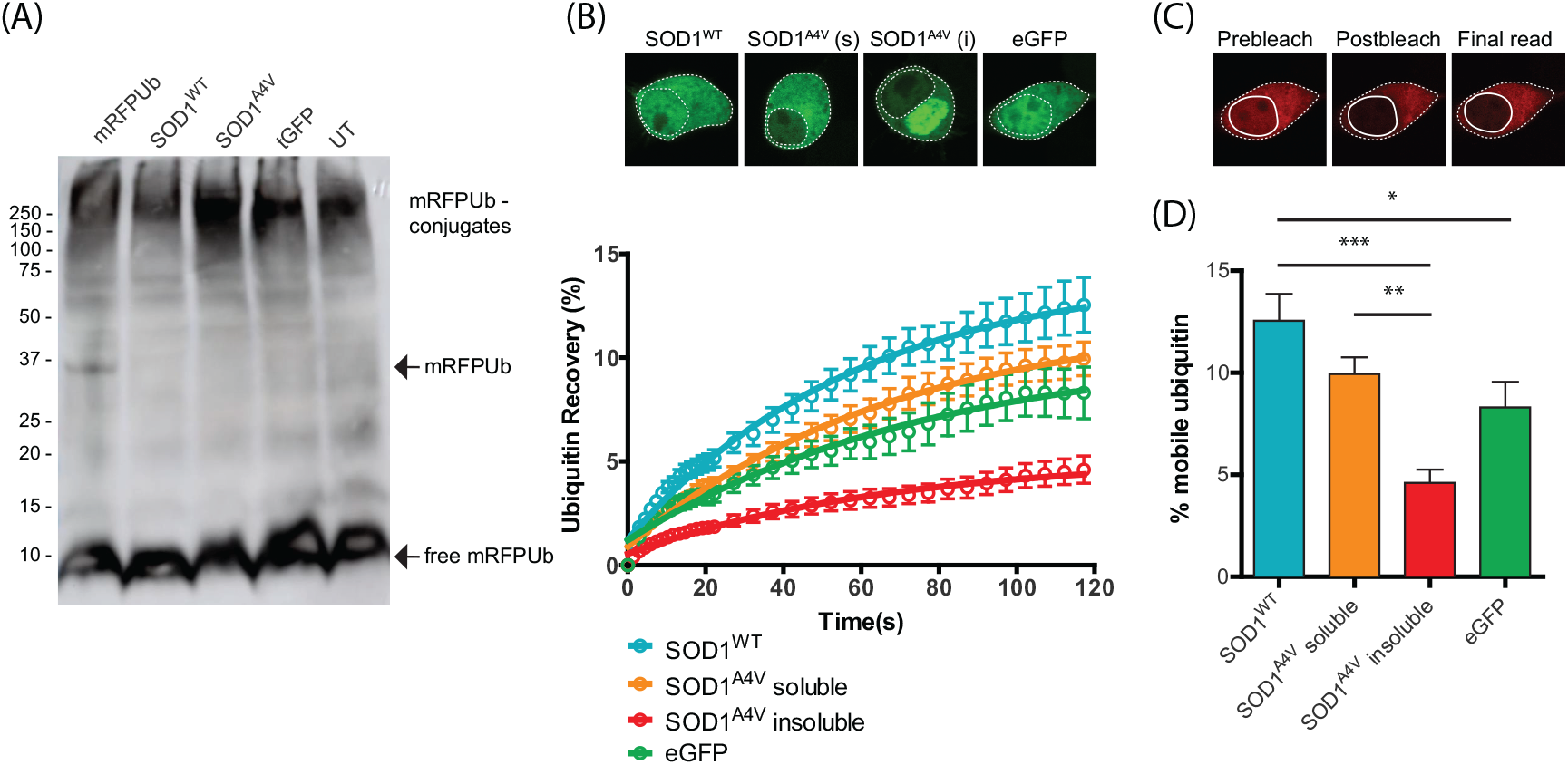
Reduced levels of free monomeric ubiquitin in NSC-34 cells containing SOD1^A4V^ aggregates. (A) Western blot analysis of cell lysates of NSC-34 cells transiently transfected with SOD1-GFP, mRFP-Ub or eGFP control. Samples were separated under reducing conditions and probed with anti-ubiquitin antibody. (B) The entire nucleus of co-transfected NSC-34 cells was photobleached and the recovery of nuclear Ub was monitored as a proportion of cytoplasmic fluorescence for 120 s. Data shown are means ± SEM (n ≥ 17) combined from 3 independent experiments. (C) Representative confocal images of pre-bleach, post-bleach and final read. (D). The percentage of mobile Ub in the nucleus at the final read was quantified as a proportion of cytoplasmic fluorescence. Data represent mean ± SEM (n ≥ 17, combined from 3 independent experiments)* P < 0.05, ** P < 0.01, *** P<0.001.

### SOD^A4V^-GFP induced changes in the ubiquitome of NSC34 cells

Free Ub exists in complex equilibrium with multiple conjugated forms and ALS mutations may induce redistribution of Ub through altered activity in various cellular pathways. To investigate changes in the Ub-modified proteome (ubiquitome) of cells expressing SOD^WT^ compared with SOD^A4V^-GFP, we performed proteomics following enrichment of cell lysates for ubiquitylated proteins (**Figure 6A**). We identified 316 ubiquitylated proteins common to cells expressing either wild-type or mutant SOD1. 55 proteins were uniquely present in the ubiquitome of cells expressing mutant SOD1 and 12 unique proteins were identified in the ubiquitome of cells expressing wild-type SOD1 (**Figure 6B, and Table 1**). Network analysis of known protein-protein interactions and ontology in the set of ubiquitylated proteins unique to cell expressing mutant SOD1 using the String database [42] identified a number of enriched pathways, including metabolic pathways, ubiquitin-proteasome system (UPS), ribosome and mRNA processing and transport (**Figure 6C**). Expression of a chronic misfolded protein has previously been observed to increase proteome misfolding and we have shown that ALS aggregates are composed of supersaturated proteins - that is, proteins which have expression levels higher than one might predict given their solubility [5].

**Figure 6:**
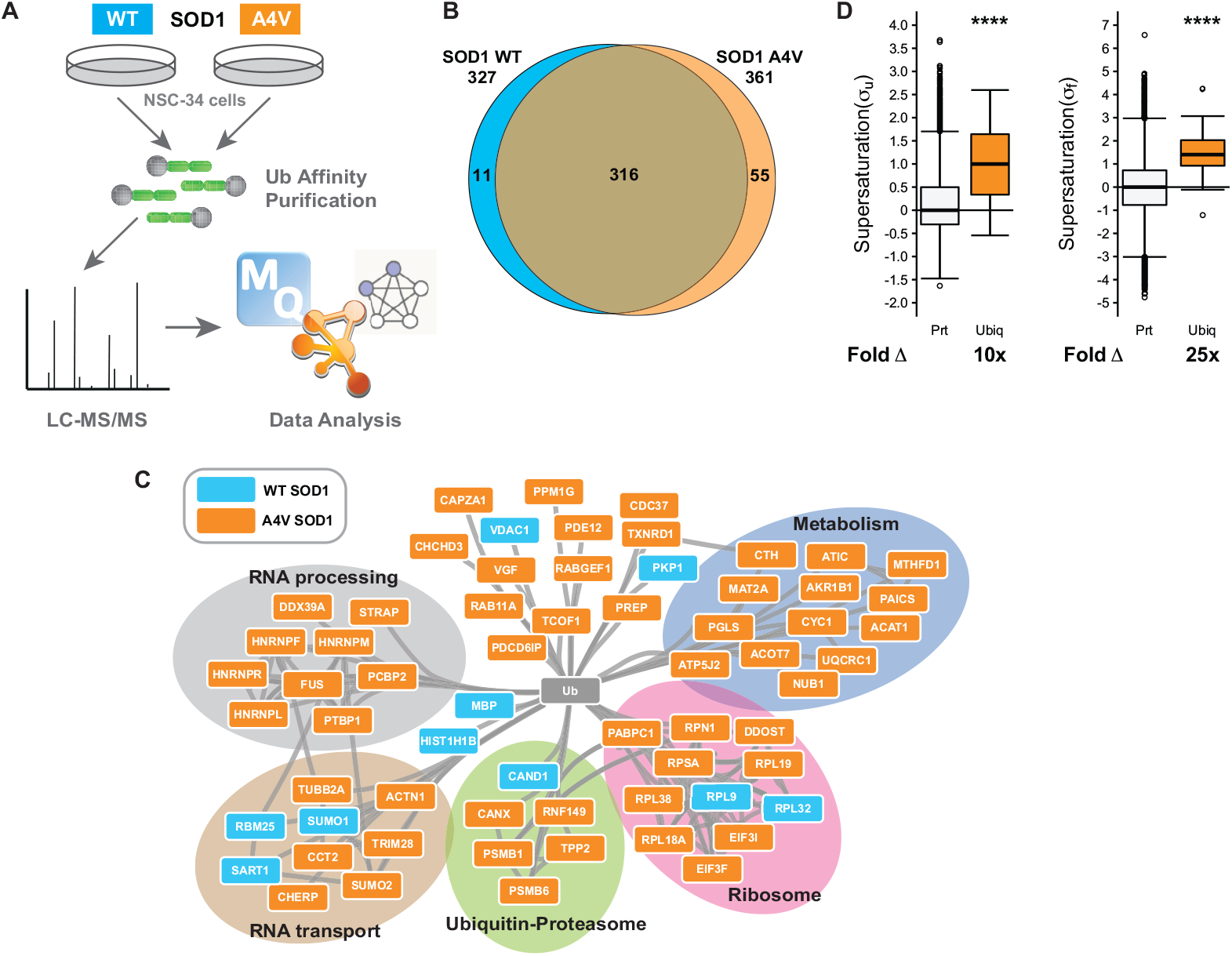
The ubiquitylated proteome of transfected NSC-34 cells. (A) NSC-34 cells expressing GFP fusions of SOD1^WT^ or SOD1^A4V^ were subjected to high affinity purification to isolate ubiquitylated proteins which were subsequently identified by LC-MS/MS. (B) Venn analysis comparing the number of ubiquitylated proteins in NSC-34 cells transfected with either SOD1^A4V^ (n=2) or SOD1^WT^ (n=3). (C) STRING analysis of protein-protein interaction network of ubiquitylated proteins unique to either SOD1^A4V^ or SOD1^WT^. Proteins were identified as present in each condition if they were present in at least two replicates for that condition. Additionally proteins were only considered “unique” to either SOD1^WT^ or SOD1^A4V^ if they were not identified in any of the replicates for any of the other conditions. (D) The median supersaturation score calculated for the unfolded (**σ**u) and native (**σ**f) states of proteins unique to the SOD1^A4V^ ubiquitome. Fold **Δ** refers to the increase in supersaturation score from the known mouse proteome.

**Table 1.**
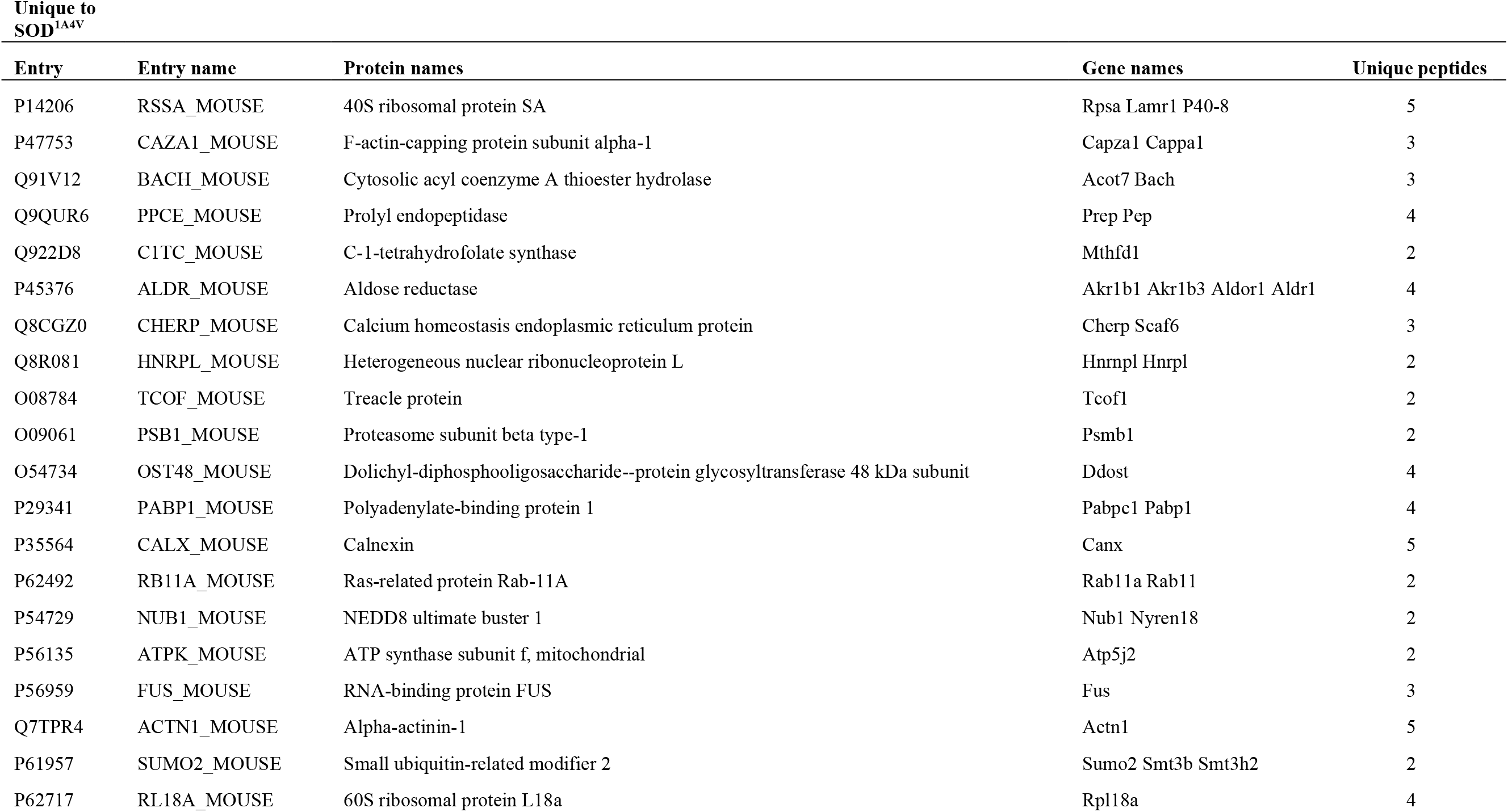
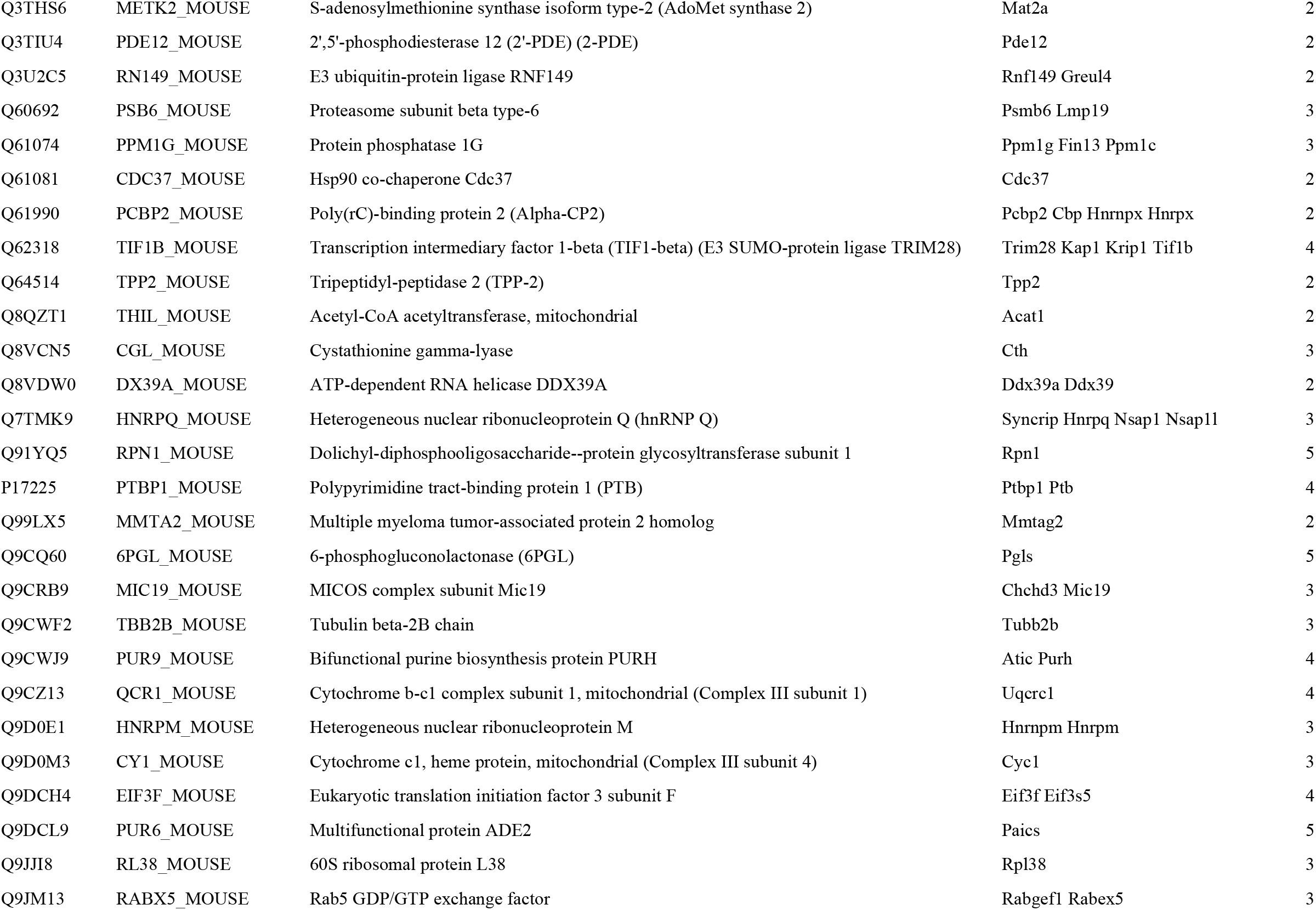
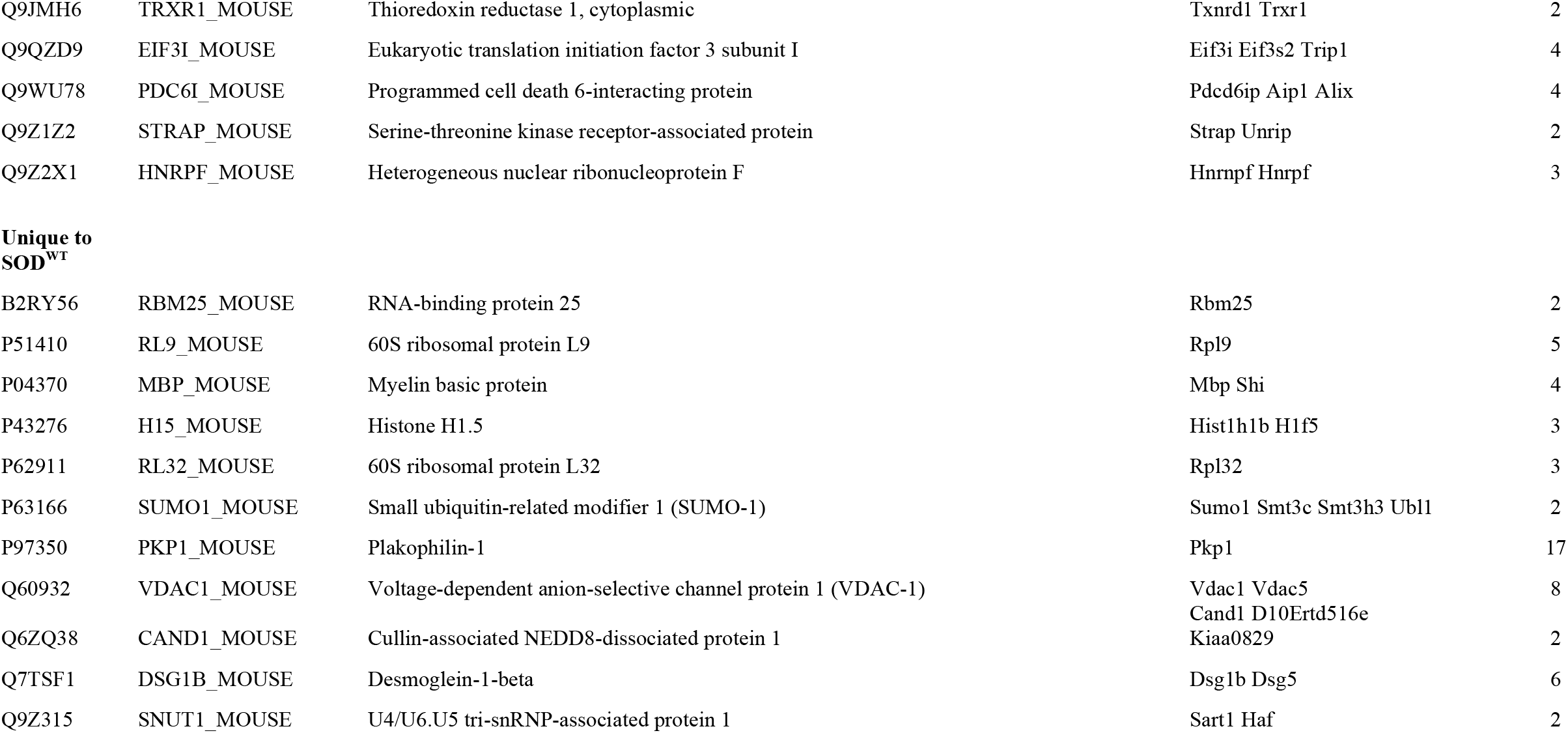
Proteins identified unique to A4V or WT SOD1:

We compared the supersaturation scores for both the unfolded and native states of ubiquitylated proteins found uniquely in SOD^A4V^-GFP expressing cells to the whole proteome. The median supersaturation score of these 55 ubiquitylated proteins is 10 × higher than the whole proteome in the unfolded state (**σ**u) and 25 x higher than the whole proteome in the native state (**σ**f) (**Figure 6D**). These data are consistent with proteome instability and resulting redeployment of Ub driving altered Ub distribution and subsequent impairment of Ub homeostasis upon expression of a chronically misfolded protein.

### Cells with SOD1 aggregates have altered mitochondrial morphology and function

Pathway analysis of our ubiquitome data (above) indicated specific enrichment of mitochondrial/metabolic proteins in the ubiquitome of SOD1^A4V^-expressing cells (**Table 1**), suggesting that Ub redistribution/homeostasis might be associated with compromised mitochondrial function in these cells. Indeed, previous work has shown that mutations in SOD1 lead to mitochondrial dysfunction [47–49]. In order to assess mitochondrial function in SOD1A4V-expressing cells, we first examined mitochondrial morphology using Mitotracker and confocal microscopy (**Figure 7A-C**). We found that SOD1^A4V^-expressing cells contained a significantly higher number of mitochondria per cell compared to controls (**Figure 7B**). In addition, we these mitochondria are significantly more circular than those in control cells (**Figure 7C**). We quantified mitotracker fluorescence in cells using flow cytometry, as above focusing on the top 10% most fluorescent cells (based on SOD-GFP expression) to enrich for cells with aggregates (i.e. when expressing SOD1^A4V^). This subset of SOD1^A4V^-expressing cells has higher mitotracker fluorescence compared to SOD1^WT^ expressing cells exhibiting the same level of fluorescence -consistent with accumulation of mitochondria with SOD1^A4V^ expression (**Figure 7D**). In the same subset of aggregate-containing cells we observed increased fluorescence of the functional reporter CMXRos, suggesting alterations in mitochondrial membrane potential (**Figure 7E**).

**Figure 7:**
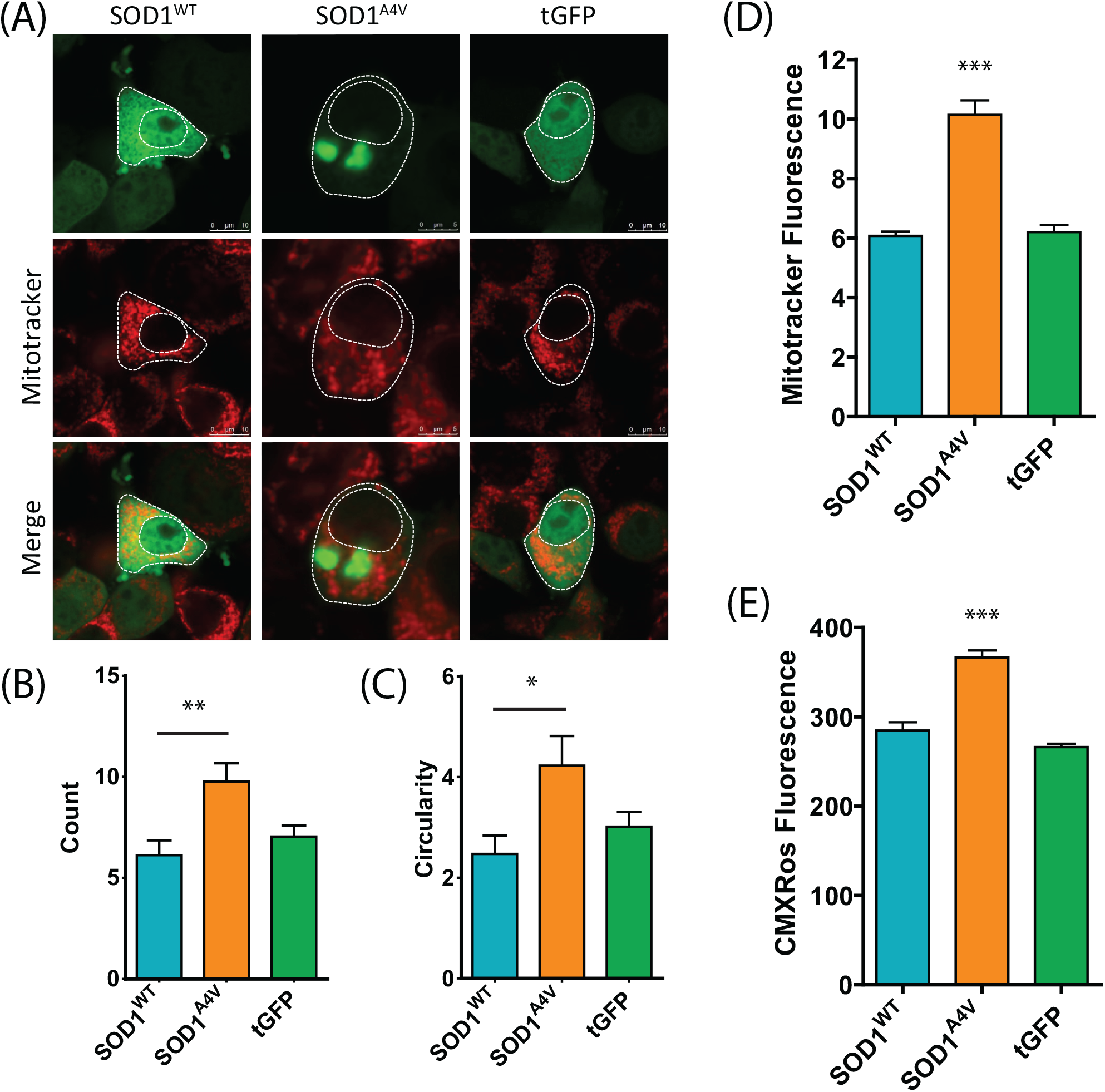
Cells containing SOD1 aggregates display altered mitochondrial morphology and dysfunction. (A) NSC-34 cells transiently transfected with SOD1-GFP were stained for mitochondria 48 h post transfection with Mitotracker Deep Red. The number of mitochondria (B) and circularity (C) was determined using a mitochondrial morphology macro in ImageJ. Data represent mean ± SEM (n ≥ 23) * P < 0.05, ** P < 0.01. (D) Mitotracker fluorescence was also quantified by flow cytometry in cells expressing the highest levels of SOD1-GFP. (E) Mitochondrial membrane potential was examined through the accumulation of Mitotracker Red CMXRos. Data represent mean ± SEM (n = 3) *** P<0.001.

## Discussion

A unifying feature of neurodegenerative diseases such as ALS is the presence of Ub within insoluble protein aggregates [10, 50, 51]. Beyond labelling substrates for degradation via the proteasome, Ub is an important regulator of cellular processes such as transcription, translation, endocytosis and DNA repair. The sequestration of Ub into inclusions may therefore reduce the availability of free Ub essential for these processes, compromising cellular function and survival. To gain insight into the regulation of Ub homeostasis in ALS, we followed the dynamic distribution of Ub at a single cell level in a well-established SOD1 cell model of ALS. Our results confirm that the expression of mutant SOD1 leads to UPS dysfunction and corresponding disruption of Ub homeostasis, suggesting these processes plays a key role in the development of ALS pathology.

In cell models, Ub regulates the concentrations of TDP-43 and SOD1 [52, 53], and proteasome inhibition can trigger their abnormal accumulation. While our previous work established that SOD1 aggregates contain Ub [45], results here confirm that SOD1 co-aggregates with Ub, and that Ub is present at the earliest stage of aggregation. Furthermore, SOD1 aggregates were found to contain both K48 and K63 polyubiquitin chains, signalling degradation via the proteasome and autophagy pathways, respectively. Interestingly, both K48 and K63-linked ubiquitylation has been associated with the formation of inclusions [54], with previous studies showing that K63 polyubiquitination directs misfolded SOD1 to the ubiquitin-aggresome route when the UPS is inhibited [55]. These data are consistent with the need for tightly regulated Ub homeostasis in neurons, and suggest any perturbation in this homeostasis could cause Ub depletion and subsequent toxicity. It is likely that protein aggregation causes Ub depletion, which could then drive further aggregation in a positive feedback loop (**Figure 8**).

**Figure 8:**
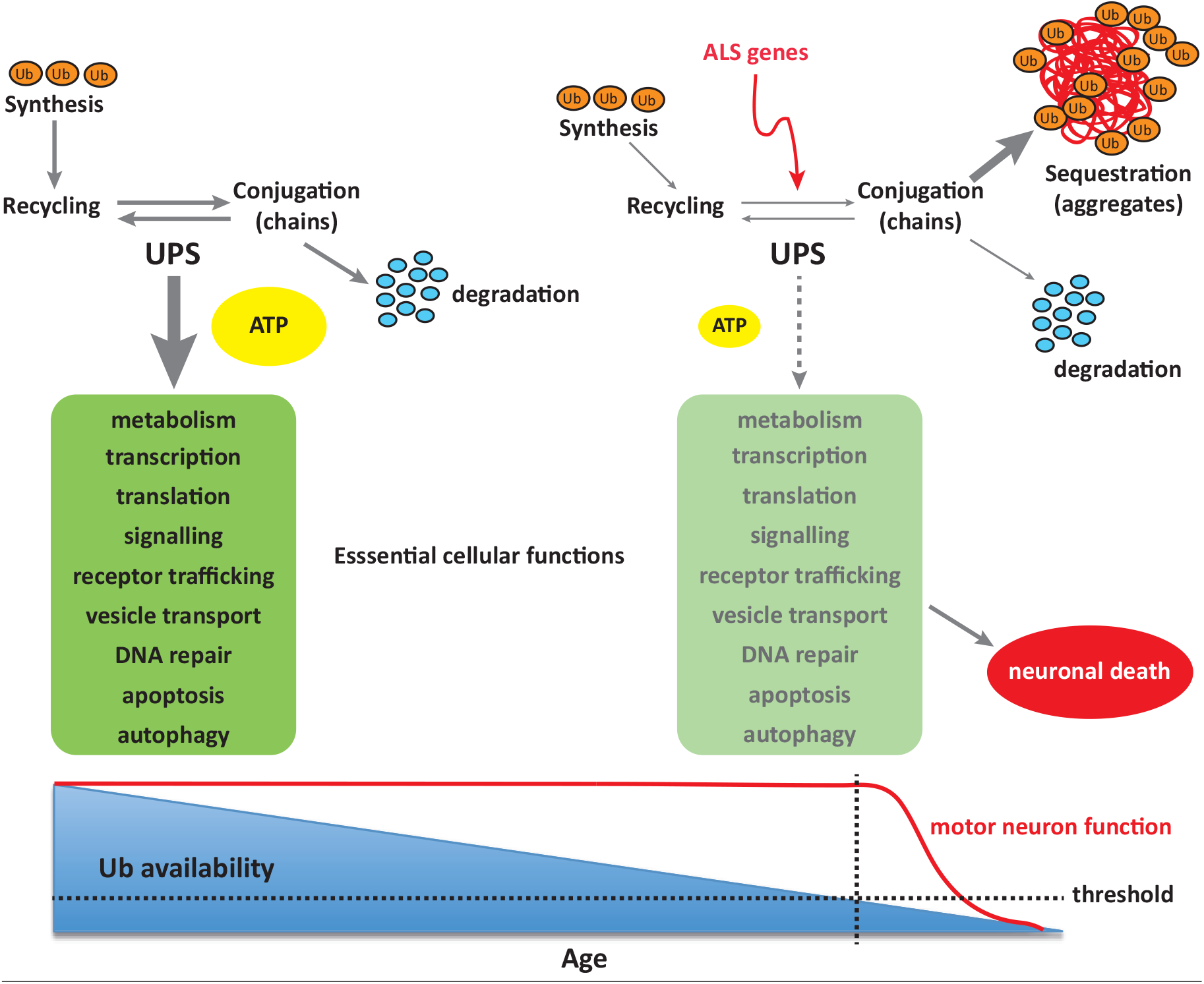
Disrupted Ub homeostasis in ALS. Mutations in SOD1-associated ALS disrupt Ub homeostasis, either directly or through sequestration of Ub into protein aggregates. These changes result in altered Ub distribution and subsequent depletion of free Ub, eventually reaching a threshold below which vital cellular functions are severely compromised, and ultimately result in cell death.

In the nervous system, the UPS contributes to the regulation of many aspects of synaptic function, such as neuronal growth and development, neuronal excitability, neurotransmission, long-term potentiation (LTP) and synapse formation and elimination [56, 57]. UPS dysfunction is therefore central to neuronal health and neurodegenerative disease [6]. Impairment of the UPS has been implicated strongly in the pathogenesis of ALS [52, 58, 59]. Here, we reveal that the aggregation of SOD1 compromises UPS function, decreasing cellular capacity to degrade the proteasome reporter tdTomato^CL1^. Our findings are consistent with the previous demonstration of UPS dysfunction in ALS, with accumulation of the fluorescent UPS reporter Ub^G76V^-GFP observed in the spinal cord and cranial motor neurons of SOD1^G93A^ mice [59]. Moreover, a motor neuron specific knockdown of proteasome subunit Rpt3 in the absence of ALS genetic background results in an ALS phenotype in mice - including locomotor dysfunction, progressive motor neuron loss and mislocalization of ALS markers TDP-43, FUS, ubiquilin 2, and optineurin [58]. These data support a model where a reduction in UPS capacity is sufficient to drive ALS pathology in mice. Not even overexpression of human mutant TDP-43 gives such an accurate reproduction of a human ALS-like phenotype in mice. In ALS patient spinal motor neurons, 85% (38/40) of cytoplasmic TDP-43 foci or inclusions were positive for Ub. In fact, almost all cellular Ub in these neurons was sequestered within large, skein-like inclusions [45]. Interestingly, Ub has been found to accumulate in inclusions without the aggregation of TDP-43 in sporadic ALS [7], suggesting that aggregation of proteins such as TDP-43, FUS and SOD1 may not be necessary for Ub depletion-induced toxicity. In human skin fibroblasts it has been shown that the UPS reporter GFP^CL1^ accumulates significantly more in ALS patient fibroblasts compared to controls, suggesting a compromised UPS [36]. This raises the question of whether there are alternate means to control protein concentration when protein aggregates disrupt degradation mechanisms. It was recently proposed that widespread transcriptional repression may serve this purpose in Alzheimer’s disease [60].

The work presented here suggests that the aggregation of misfolded SOD1 alters Ub homeostasis and subsequently depletes the free Ub pool in cells. Cellular Ub exists in a complex equilibrium between free and conjugated Ub [31]. Neurons are vulnerable to free Ub deficiency, which if prolonged may lead to cell death [61, 62]. Many factors can influence Ub homeostasis. For example, proteasome inhibition depletes free Ub to as low as 5% of basal levels in less than 2 h [63, 64]. Inhibition of translation also depletes free Ub through reduced production, while toxicity can be rescued by overexpression of Ub [65]. Accumulation of ubiquitylated proteins in inclusions is an important potential mechanism for depletion of free Ub, and in this context free Ub levels can be partially restored by overexpression or removal of Ub from the aggregated protein through ubiquitin-specific proteases. It was recently shown that while the ataxia associated mutation in Usp14 causes reduction in free Ub and neuromuscular junction dysfunction in mice, overexpression of Ub restored free Ub levels in motor neurons and improved NMJ structure [66]. Additionally, free Ub could be increased in cells containing huntingtin aggregates by overexpression of the de-ubiquitylation enzyme USP14 [67]. Not only did this protect cells from aggregate-induced toxicity, it also reduced ER stress, which is thought to precede inclusion formation in ALS models [18]. These data further support the concept that modulation of cellular Ub pools is an important factor in the pathogenesis of neurodegenerative disease.

The pathways responsible for modulating Ub homeostasis in ALS are not yet well understood. Our ubiquitomics analysis of cells expressing SOD1 reveals that the unique ubiquitome of cells expressing mutant SOD1 is enriched for proteins prone to aggregation (i.e. supersaturated). This is consistent with findings that proteins associated with neurodegeneration are supersaturated [5, 25, 26]. Furthermore, a substantial proportion of the ubiquitylated proteins identified are from pathways known to be dysfunctional in ALS, including RNA processing, metabolic and mitochondrial pathways. This is not surprising given that both RNA binding proteins and oxidative phosphorylation and metabolic pathways are also prone to aggregation in neurodegeneration [26]. These results suggest that mutations in SOD1 may lead to mitochondrial dysfunction by disrupting Ub homeostasis. Interestingly, we find that cells expressing mutant SOD1 had altered mitochondrial morphology and function. Disturbances to mitochondrial morphology have been previously reported in human motor neurons carrying the SOD1^A4V^ mutation [68] and recent studies have revealed that mitochondrial impairment occurs soon after proteasome inhibition [69]. In fact, continued proteasome dysfunction in mouse brain cortical neurons inhibited the degradation of ubiquitylated mitochondrial proteins and led to the accumulation of dysfunctional mitochondria [70]. Misfolded proteins such as mutant SOD1 have also been shown to interact with mitochondrial proteins and translocate into the mitochondrial matrix, where they accumulate and induce mitochondrial dysfunction [71–73]. However, this cytotoxicity can be attenuated through the ubiquitylation of misfolded but non-aggregated SOD1 - which promotes its degradation via the UPS [74]. Collectively, these studies demonstrate an integral functional relationship between impairment of the UPS and mitochondrial dysfunction, and that modulation of cellular Ub pools may rescue mitochondrial dysfunction caused by the accumulation of SOD1.

## Conclusions

In conclusion, we observe that the aggregation of mutant SOD1 leads to altered UPS activity and redistribution of Ub, causing disrupted Ub homeostasis. Given that Ub controls many essential cellular pathways that are also dysfunctional in ALS - including transcription, translation, vesicle transport, mitochondrial function and apoptosis - these findings suggest that Ub homeostasis is a central feature of ALS pathogenesis. Further understanding of the contribution of Ub homeostasis to ALS pathology may be imperative to understanding the molecular pathways underpinning neurodegenerative disease more broadly.

## List of abbreviations

Amyotrophic lateral sclerosis (ALS)

Bovine serum albumin (BSA)

Deubiquitinating enzymes (DUBS)

Familial ALS (fALS)

Fluorescence loss in photobleaching (FLIP)

Fluorescence recovery after photobleaching (FRAP)

Frontotemporal dementia (FTD)

Juxtanuclear quality control compartment (JUNQ)

Long-term potentiation (LTP)

Motor neuron disease (MND)

Neuromuscular junction (NMJ)

Paraformaldehyde (PFA)

Phosphate buffered saline (PBS)

Region of interest (ROI)

Room temperature (RT)

Sporadic ALS (sALS)

Superoxide dismutase 1 (SOD1)

TritonX-100 (TX-100)

Ubiquitin (Ub)

Ubiquitin-proteasome system (UPS)

## Declarations

### Ethics approval and consent to participate

Not applicable

### Consent for publication

Not applicable

### Availability of data and material

The datasets used and/or analysed during the current study available from the corresponding author on reasonable request.

## Authors’ contributions

N.E.F, P.C, L.M, K.L.V, D.N.S, and J.J.Y designed research; N.E.F, P.C, I.A.L.S, L.M, J.M, and K.M performed research; N.E.F, I.A.L.S, P.C, L.M, K.L.V, D.N.S, and J.J.Y analyzed data; and N.E.F, P.C, D.N.S, and J.J.Y wrote the paper. All authors read and approved the final manuscript.

## Acknowledgements

Not applicable

Authors’ information (optional)

## Supplementary Methods

To quantify SOD1 inclusion formation, images were acquired as described previously [1]. Briefly, z-stack images of NSC-34 cells transfected with SOD1-GFP constructs were acquired 48 h post transfection using the 63X objective, 512 x 512 pixels, 1 line and frame average and a scan speed of 400 Hz. Z-stacks were processed into a single image using LAS-AF Lite software (Leica) and at least 100 transfected cells were analysed for the presence of inclusions.

**Toxicity Assay**. The toxicity of cells expressing SOD1 was monitored over 68 h in an Incucyte automated fluorescent microscope (Essen BioScience, USA) as described in [1]. Cells were dissociated 24 h post transfection and replated in 96-well plates at a confluency of 20% in phenol red free DMEM/F12 containing 10% FBS. Images were acquired every 2 h and analysed using a processing definition trained to select GFP positive cells. The number of GFP positive cells was normalised to time zero before SOD1^A4V^ numbers were adjusted to SOD1^WT^ values.

**Figure S1:**
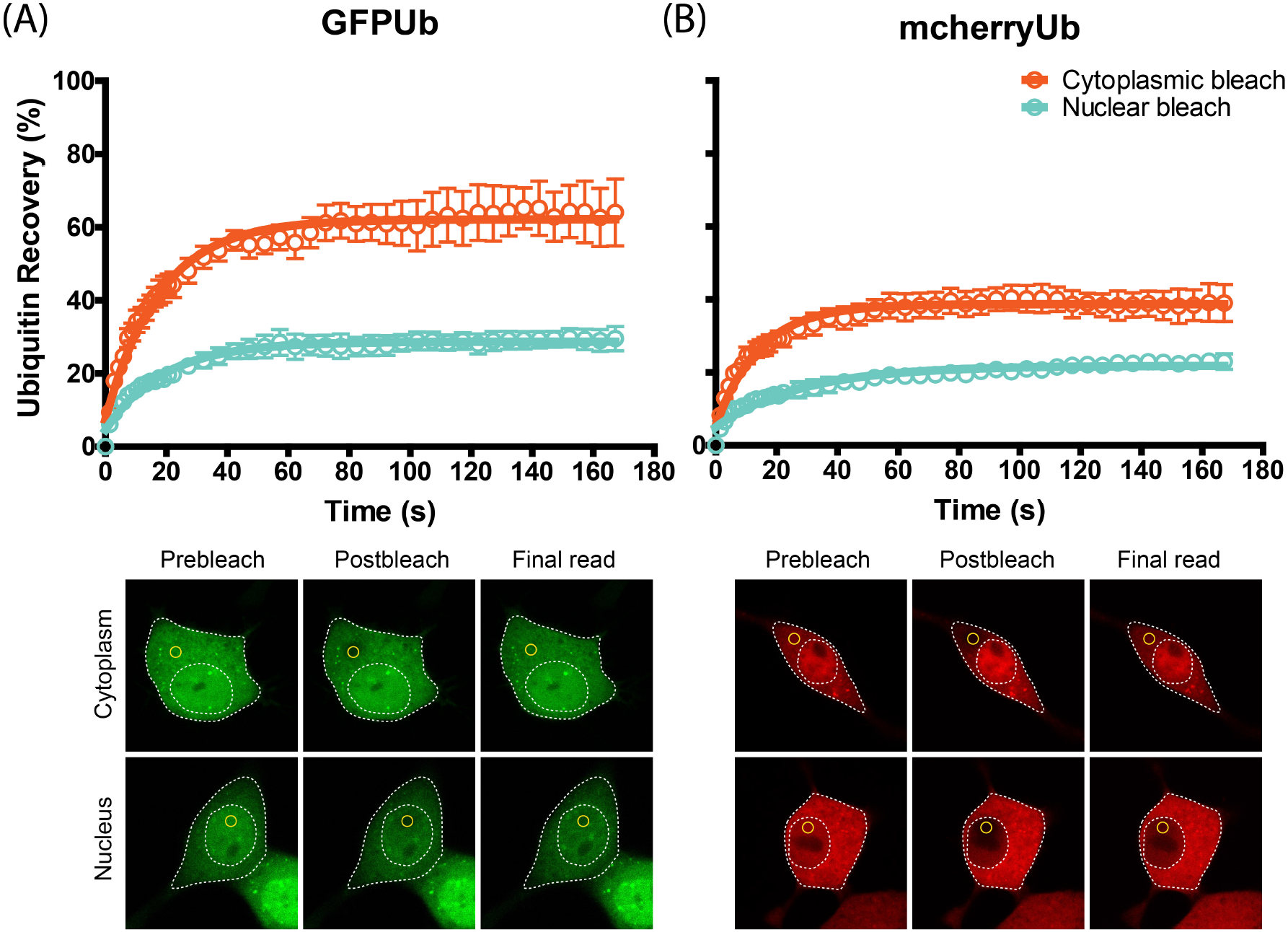
mcherry-Ub behaves in a similar manner to GFP-Ub. (A) NSC-34 cells transfected with either GFP-Ub (A) or mcherry-Ub (B) were photobleached in the cytoplasm or nucleus and recovery of Ub fluorescence was monitored for up to 170 s. Representative confocal images of prebleach, postbleach and recovery endpoint are shown with the ROI marked in yellow.

**Figure S2:**
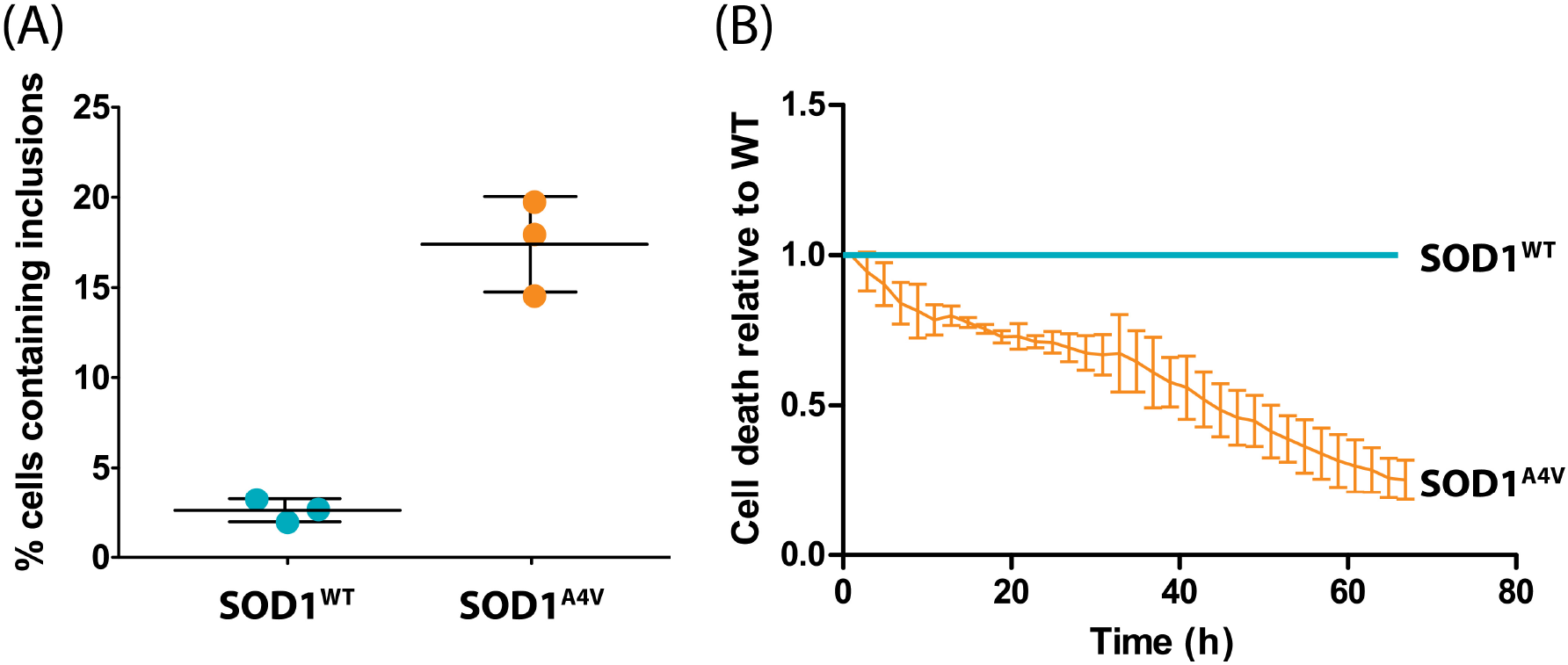
SOD1A4V expression induces inclusion formation and is associated with increased cell death in NSC-34 cells. (A) Inclusion formation in NSC-34 cells expressing SOD1-GFP was determined 48 h post transfection. (B) The toxicity of SOD1^A4V^ was monitored in comparison to SOD1WT over a 68 h period.

